# CARD8 negatively regulates NLRP1 inflammasome activation level by interaction with NLRP1

**DOI:** 10.1101/2022.06.26.497666

**Authors:** Zhihao Xu, Shasha Deng, Yuluo Huang, Yunru Yang, Liangqi Sun, Hanyuan Liu, Dan Zhao, Weihong Zeng, Xueying Yin, Peiyi Zheng, Muziying Liu, Weidong Zhao, Ying Zhou, Tengchuan Jin

**Author notes:** Corresponding author: Ying Zhou or Tengchuan Jin.

## Abstract

NLRP1 inflammasome functions as canonical cytosolic sensor in response to intracellular infections and is implicated in auto-inflammatory diseases. But the regulation and signal transduction mechanisms of NLRP1 are incompletely understood. Here, we show that the T60 variant of CARD8, but not the canonical T48 isoform, negatively regulates NLRP1 inflammasome activation by directly interacting with the receptor molecule NLRP1 and inhibiting inflammasome assembly. Furthermore, our results suggest that the different ASC preference in three types of inflammasomes, namely ASC-indispensable NLRP1 inflammasome, ASC-dispensable mNLRP1b inflammasome and ASC-independent CARD8 inflammasome, is mainly caused by the CARD domain, not the UPA subdomain. Based on the systematic site-directed mutagenesis and structural analysis, we find that the signal transduction of NLRP1 inflammasome relies on multiple interaction surfaces on its death domain superfamily member CARD domain. Finally, our results partly explain the mechanism of the NLRP1 mutation-derived auto-inflammatory diseases caused by the overactivation of the NLRP1 inflammasome. In conclusion, our study not only reveals how CARD8 downregulates NLRP1 inflammasome activation, but also provides insights into the mechanisms of CARD-containing inflammasome assembly.

## Introduction

The canonical inflammasomes are cytosolic multiprotein complexes that initiate pyroptotic cell death and inflammatory cytokine secretion in response to diverse pathogenic and endogenous danger signals^1^. NLRP1, the first discovered inflammasome sensor, could assemble inflammasome macromolecular platform by oligomerizing with the ASC (also known as PYCARD) and recruiting the effector inflammatory molecule pro-Caspase-1^2^. It facilitates the processing of pro-interleukin-1β (pro-IL-1β), pro-interleukin-18 (pro-IL-18) and gasdermin D (GSDMD) that results in releasing the maturated cytokines and inducing inflammatory cell death. Recently, some specific pathogen-associated molecular patterns (PAMPs), such as enteroviral 3C proteases and the double-stranded RNA, were identified that can activate the human NLRP1 inflammasome^3, 4^. In addition, the dipeptidyl peptidases DPP8/9 can directly interact with FIINDs domain of human NLRP1 and CARD8^5–7^. Consistently, DPP8/9 inhibitors, such as Val-boroPro (VbP), induce NLRP1 and CARD8 inflammasome assembly through their UPA-CARD domains generated upon proteasome-mediated degradation of their respective N-terminal fragments^8–11^.

In addition, the mutations on the NLRP1 result in constitutive NLRP1 inflammasome overactivation and inflammatory cytokines secretion, which have been associated with a number of chronic inflammatory disorders, including autoinflammation with arthritis and dyskeratosis (AIADK) and familial keratosis lichenoides chronica (FKLC)^12, 13^. NLRP1 inflammasome needs to keep in the inactive state to prevent aberrant activation and damaging inflammation in absence of infection and danger related molecules. Thus, understanding the negative regulation mechanism of NLRP1 inflammasome signaling is critical for discovery of novel therapy strategy for these auto-inflammatory diseases.

It is known that NLRP1 and CARD8 share similar domain organization, which both possess C-terminal function-to-find domain (FIIND) and caspase activation and recruitment domain (CARD). The FIIND domain has a proteolytic site to split into ZU5 (found in ZO-1 and UNC5) and UPA (conserved in UNC5, PIDD, and ankyrins) subdomains with a noncovalent association for the auto-inhibited N-terminal fragments. For CARD domain, it is a member of death domain (DD) superfamily that engage in assembly of inflammasome complex by homotypic and heterotypic interactions. The detailed description for downstream molecules recruitment of CARD-containing sensor proteins has not been presented. In addition, it has been reported that CARD8 plays a role as a negative regulator in humans, but not in rodents. Some investigators showed that CARD8 binds to NOD2 and suppresses NOD2-dependent inflammatory signal transduction^14^. What’s more, recent publications have reported that CARD8 interacts physically with NLRP3 to prevent its binding to the adaptor protein ASC during NLRP3 inflammasome activation^15, 16^. Thus, more studies are needed to unravel the association of CARD8 with NLRP1 based on structural homology of NLRP1 and NLRP3. In this study, we not only identify that CARD8 negatively regulates NLRP1 inflammasome activation through its direct interaction with NLRP1, but also provide structural and molecular basis for the specificity of CARD-mediated inflammasomes assembly.

## Results

### CARD8-T48 isoform has an interaction with NLRP1 but fails to inhibit its functions

Previous studies have demonstrated that CARD8 is not only an inflammasome-forming sensor, but also acts as a negative regulator in several pro-inflammatory and apoptotic signaling pathways^16–, 18^. To further evaluate the exact role of CARD8 in NOD-like receptors family, we first predicted the protein-protein interaction network with an analysis tool STRING. Despite NLRP1 and CARD8 having a structural homology, the molecular association of CARD8 and NLRP1 remains to be clarified and verified experimentally (Fig. S1a). Then, we examined the tissue expression profile of multiple inflammasome components using available 27 different human organs and tissues transcriptome data^19^. Based on specimens from altogether 95 individuals, we found that CARD8 and NLRP1 are widely expressed, in contrast to other known inflammasome receptors (Fig. 1a). In addition, CARD8 possesses a similar tissue expression profile as compared to NLRP1, perhaps indicating that CARD8 and NLRP1 have a functional molecular association. Based on these observations, the analysis of the CARD8 polymorphism was performed to investigate the regulatory role of CARD8 in inflammasome signal transduction pathways. CARD8 mainly contains six different isoforms, among which the T48 isoform was considered as canonical sequence previously (Fig. 1b). Compared with the T48 isoform, the T60 isoform exhibited a longer unstructured N-terminal region and an additional alanine in the ZU5 subdomain. This may show that distinct isoforms of CARD8 play different roles in mediating signal transduction, and the functional differences among CARD8 isoforms are unknown. To investigate the interaction between wild-type full-length CARD8 and NLRP1, co-immunoprecipitation assay was performed. And the result showed that CARD8-T48 isoform was immunoprecipitated with NLRP1, suggesting that CARD8-T48 isoform can bind to NLRP1 in HEK 293T cells (Fig. 1c). To ascertain whether CARD8 could affect the NLRP1 inflammasome activation, we co-transfected NLRP1 inflammasome components to reconstitute the NLRP1 inflammasome in HEK 293T cells. The co-expression of NLRP1, ASC, pro-Caspase-1 and pro-IL-1β can result in the caspase-1-mediated IL-1β maturation and secretion, showing that the NLRP1 reconstitution system could be used to study the NLRP1 inflammasome regulation mechanism (Fig. 1d and e). Then, the CARD8-T48 isoform was introduced into HEK 293T cells to determine the regulatory function of CARD8 using the reconstitution system. We found that CARD8-T48 isoform did not cause a defect in processing of pro-IL-1β by Caspase-1, indicating no inhibitory function of CARD8-T48 on NLRP1 inflammasome activation in HEK 293T cells (Fig. 1f and g). Taken together, these results showed that CARD8-T48 isoform binds with NLRP1 but fails to inhibit NLRP1 inflammasome activation.

**Fig. 1.**
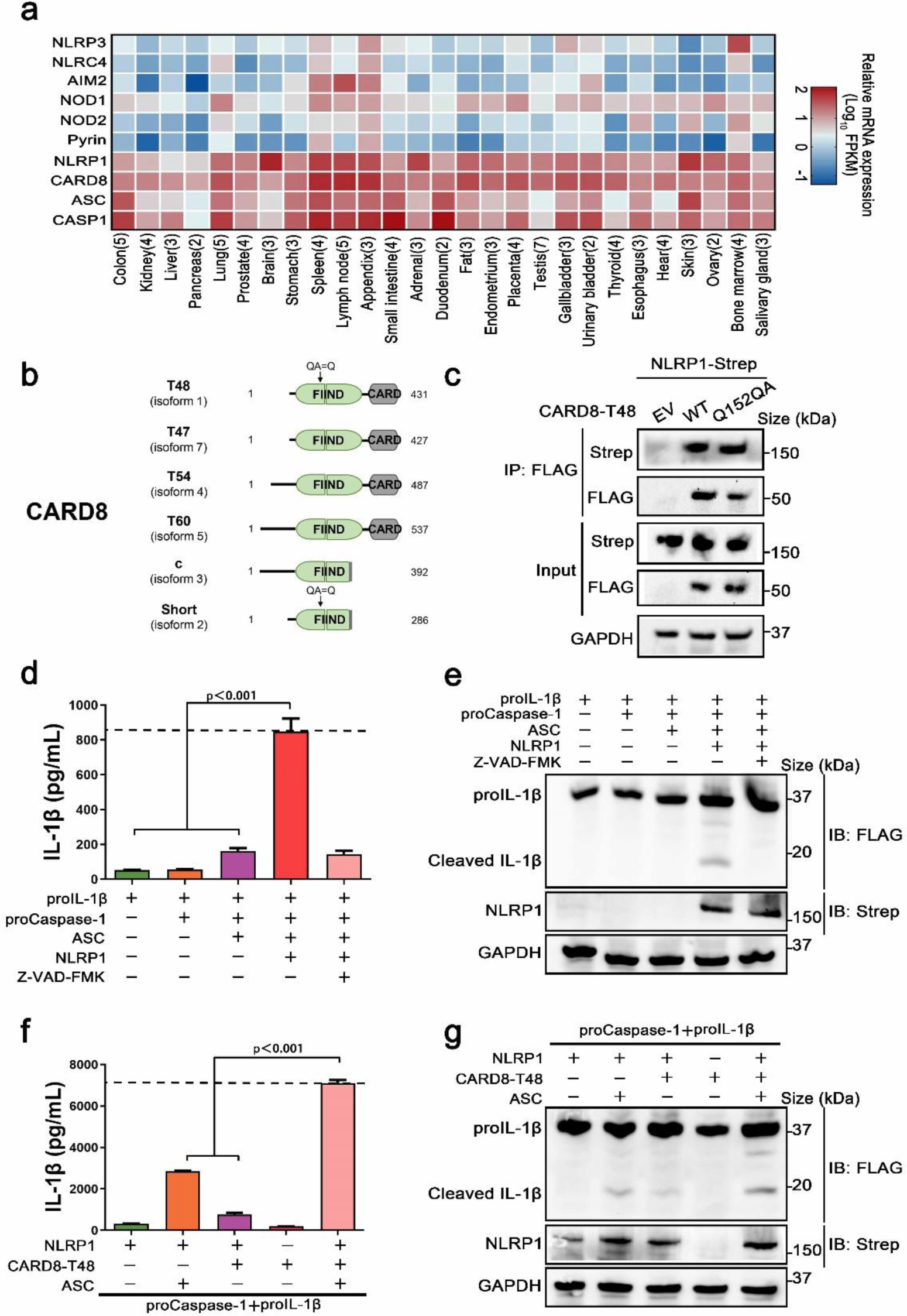
CARD8-T48 isoform interacts with NLRP1, but it does not inhibit NLRP1 activation. **a** Heat map of NLRP1 and other inflammasome sensors transcription expression levels in 27 kinds of tissues or organs in human body. The numbers of replicate samples are labeled in parentheses. **B** Diagram of human CARD8 different isoforms. Full-length CARD8 protein consists of a variable aminoterminal disordered region, FIIND and CARD domain. **c** HEK 293T cells were transiently transfected with constructs for empty vector (EV), CARD8-T48, CARD8-T48-Q152QA and NLRP1. After 28 hours, the protein expression was detected by immunoblotting, and co-immunoprecipitation (IP) were conducted with anti-FLAG antibody to analyze the interaction. **d,e** HEK 293T cells were transiently transfected with constructs encoding the pro-IL-1β, proCaspase-1, ASC and NLRP1. At 48 hours later, the cultural supernatants were monitored for cleaved IL-1β by ELISA and cell lysates were evaluated by immunoblotting. Data are shown as mean ± SEM and representative of three biological replicates. **f,g** HEK 293T cells were transiently transfected with constructs encoding the pro-IL-1β, proCaspase-1, ASC, NLRP1 and CARD8-T48. After 48 hours, ELISA and immunoblotting analysis were conducted with cell cultural supernatants and lysates. Data are shown as mean ± SEM and representative of three biological replicates.

### Negative regulation of NLRP1 inflammasome activation level by CARD8-T60 isoform

As noted above, the CARD8 has six different isoforms and similar tissue distributions. To identify whether other different CARD8 isoforms could affect the NLRP1 inflammasome activation, we first characterized the expression levels of distinct CARD8 isoforms in 27 different human organs and tissues. To our surprise, CARD8-T60 isoform exhibited a much higher expression level compared with other CARD8 isoforms (Fig. 2a). To investigate the effect of CARD8-T60 isoform in NLRP1 inflammasome, CARD8-T60 isoform was expressed, which leading to significantly decreased cleavage products of pro-IL-1β (Fig. 2b and c). To further examine the influence of CARD8-T60 on NLRP1-mediated activation, the maturation and secretion level of IL-1β in the expression of different dosages of CARD8-T60 was measured. In the reconstituted NLRP1 inflammasome system, the secretion level of Caspase-1-mediated release of IL-1β was considerably reduced in a dose-dependent manner (Fig. 2d and e).

**Fig. 2.**
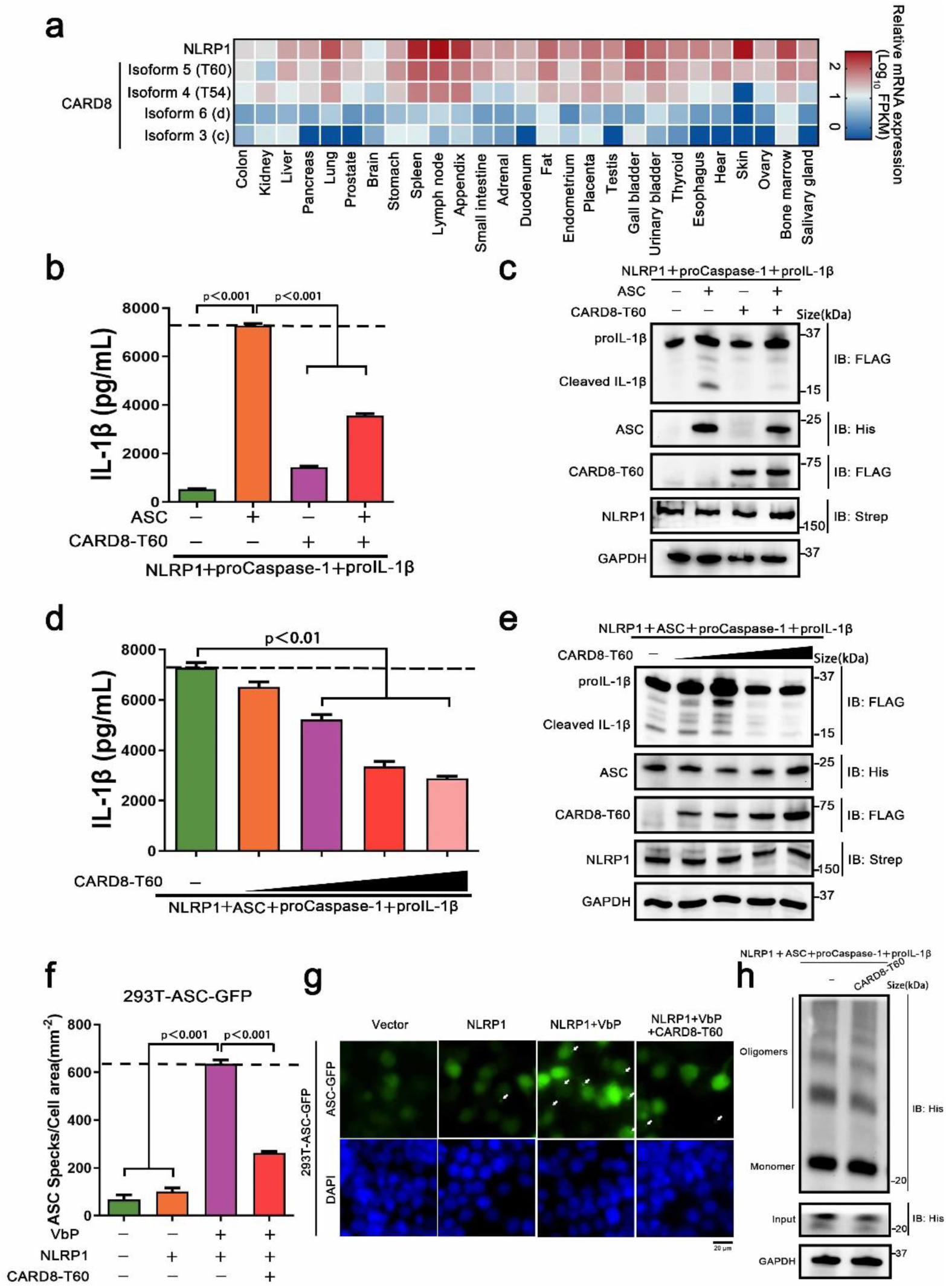
CARD8-T60 isoform suppresses NLRP1 inflammasome activation level. **a** Heat map of NLRP1 and CARD8 isoforms transcription expression levels in 27 kinds of tissues or organs in human body. **b,c** HEK 293T cells were transiently transfected with constructs encoding the pro-IL-1β, proCaspase-1, ASC, NLRP1 and CARD8-T60. After 48 hours, ELISA and immunoblotting analysis were conducted with cell cultural supernatants and lysates. Data are shown as mean ± SEM and representative of three biological replicates. **d,e** HEK 293T cells were transiently transfected with pro-IL-1β, proCaspase-1, ASC, NLRP1 and an increased concentration CARD8-T60 expression plasmids. After 48 hours, ELISA and immunoblotting analysis were conducted with cell cultural supernatants and lysates. Data are shown as mean ± SEM and representative of three biological replicates. **f,g** HEK 293T cells stably expressing ASC-GFP were transiently transfected with the indicated expression plasmids and treated with VbP (5 μM) for 6 hours. The cells were fixed with 4% formaldehyde and evaluated for ASC speck formation by fluorescence microscopy in three differential regions. Data of mean specks per cell area are shown as mean ± SEM in f and representative images are shown in g. **h** HEK 293T cells were transiently transfected with constructs encoding the pro-IL-1β, proCaspase-1, ASC, NLRP1 and CARD8-T60. Cell lysates were subjected to crosslink with 2 mM DSS and visualized by immunoblotting.

We sought to investigate the molecular mechanism of CARD8-T60 isoform on NLRP1 inflammasome activation. As inflammasome signal platform assembly is essential for NLRP1 inflammasome activation, we next corroborated the role of CARD8-T60 in the assembly of NLRP1 inflammasome. HEK 293T cells were stably expressed with ASC-GFP fluorescent fusion protein to examine the number of ASC-included speck-like aggregates (Fig. S1b). By detection of ASC-speck formation in 293T-ASC-GFP cells, we found that NLRP1 inflammasome assembly was significantly diminished in the presence of the full-length CARD8-T60 isoform (Fig. 2f and g). Consistent with the microscopy results, we cross-linked cell lysates that overexpressed NLRP1 inflammasome components and found that ASC oligomerization was abrogated in expression of CARD8-T60 isoform (Fig. 2h). Altogether, these findings showed that CARD8-T60 isoform could suppress the formation and activation of NLRP1 inflammasome in a dose-dependent manner.

### CARD8-T60 directly interacts with NLRP1

In order to clarify the molecular mechanism of CARD8-T60 isoform inhibits NLRP1 inflammasome activation, we next analyzed the interaction between CARD8-T60 and the components of NLRP1 inflammasome. Co-immunoprecipitation was confirmed between CARD8-T60 and full-length NLRP1, but not ASC or pro-Caspase-1 (Fig. 3a). With the co-expression of RFP-CARD8 and NLRP1-GFP, the fluorescent signal showed significant co-localization of CARD8 and NLRP1 in HEK 293T cells (Fig. 3b). The similar cytoplasmic distribution of CARD8 and NLRP1 suggests that CARD8 and NLRP1 interact with each other in cells. Furthermore, we observed that RFP-CARD8 was also colocalized with murine mNLRP1b in a high colocalization coefficient as with human NLRP1, suggesting that CARD8-NLRP1 interaction is an evolutionarily conserved event in NLRP1 inflammasome (Fig. S1c and d). To rule out other molecules mediated the interaction of CARD8 and NLRP1, we evaluated the interaction of CARD8 and NLRP1 using purified recombinant CARD8 and NLRP1 proteins in an *in vitro* proteins solution (Fig. S1e). Soluble Flag-tagged NLRP1 and CARD8 proteins were purified using mammalian protein expression system (Fig. 3c). Further co-immunoprecipitation assay showed that NLRP1 was pulled down by CARD8 protein which suggest their direct interaction (Fig.3d).

**Fig. 3.**
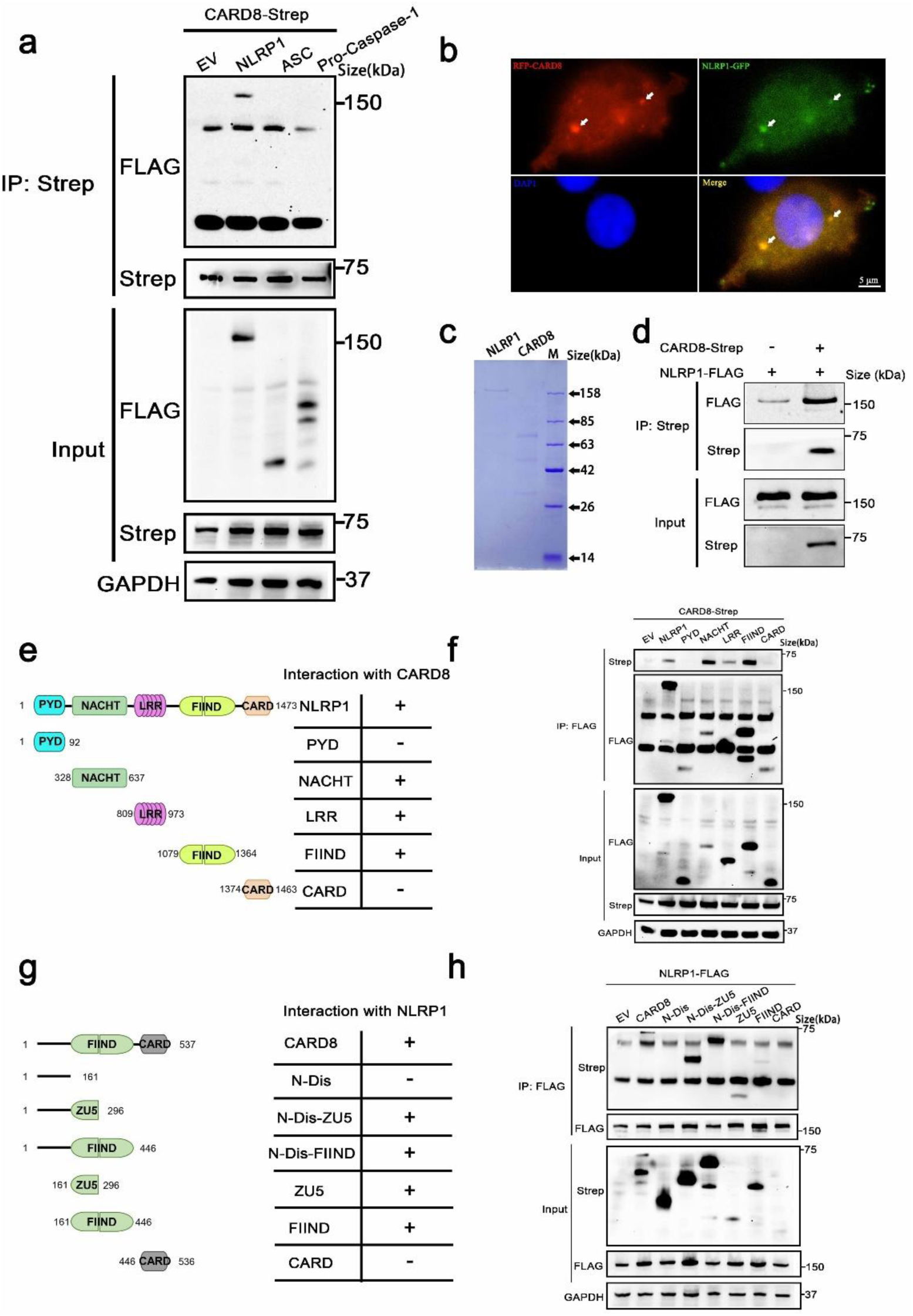
CARD8-T60 co-localizes and directly interacts with NLRP1. **a** HEK 293T cells were transiently transfected with constructs for empty vector (EV), NLRP1, ASC, proCaspase-1and CARD8-T60. After 28 hours, the protein expression was detected by immunoblotting, and co-immunoprecipitation were conducted with anti-Strep antibody to analyze the interaction. **b** Transient transfection RFP-CARD8 and NLRP1-GFP in HEK 293T cells. Green and red fluorescent were captured by fluorescence microscopy. Co-localization regions are labeled with white arrow. **c** The SDS-PAGE analysis of recombinant NLRP1 and CARD8 proteins. **d** Pull down assay of purified NLRP1-FLAG and CARD8-Strep proteins *in vitro*. **e** Diagrammatic representation of truncation mutants of NLRP1. **f** HEK 293T cells were transiently transfected with constructs for empty vector (EV), NLRP1, NLRP1 truncation mutants and CARD8-T60. After 28 hours, the protein expression was detected by immunoblotting, and co-immunoprecipitation were conducted with anti-FLAG antibody to analyze the interaction. **g** Diagrammatic representation of truncation mutants of CARD8-T60. **h** HEK 293T cells were transiently transfected with constructs for empty vector (EV), CARD8, CARD8 truncation mutants and NLRP1. After 28 hours, the protein expression was detected by immunoblotting, and co-immunoprecipitation were conducted with anti-FLAG antibody to analyze the interaction.

Moreover, to ascertain the details of CARD8-NLRP1 interaction, we conducted a systematic mapping analysis of binding of CARD8 with NLRP1. The co-immunoprecipitation experiments were performed using different domains of NLRP1 with the full-length CARD8 and vice versa. This analysis revealed that the CARD8-binding domains in NLRP1 are NACHT, LRR and FIIND domains (Fig. 3e and f). Similarly, we found that CARD8-NLRP1 interaction required the minimal ZU5 subdomain of CARD8 (Fig. 3g and h). Comparing the molecular structure of the T48 and T60 isoforms, T60 has a longer N-terminal disordered region and an additional alanine at the ZU5 subdomain. And this difference results in a completely different regulation on the NLRP1 inflammasome. The presence of N-terminal unstructured region of the CARD8-T60 isoform could increase the binding capability between NLRP1 and CARD8, but the mutant of CARD8-T48 isoform with an alanine inserted in its ZU5 subdomain (Q152QA) has no significant change in the CARD8-T48 and NLRP1 interaction (Fig. 1c and 3h). These results demonstrated once again that CARD8 directly binds NLRP1 to inhibit its activation level and there is a constitutive interaction of CARD8 with NLRP1 where the ZU5 (CARD8) subdomain binds to the NACHT, LRR, FIIND domain of NLRP1.

### Distinct ASC preference is dependent on CARD, but not UPA domain, in NLRP1, mNLRP1b and CARD8 inflammasome

During the investigation into the effect of CARD8 on NLRP1 inflammasome, the data show that human CARD8 could induce the secretion of IL-1β in the absence of ASC, while NLRP1 does not (Fig. 1f and 2b). The inflammasome assembly is extremely complicated and little is known about the assembly mechanism of CARD-containing inflammasomes, including mouse mNLRP1b, human NLRP1 and CARD8 inflammasome. We aim to systematically analyze the ASC preference in these CARD-containing inflammasomes. The adaptor protein ASC is necessary for IL-1β secretion in human NLRP1 inflammasome, while no significant differences in IL-1β secretion were observed in the presence or absence of ASC for human CARD8 inflammasome (Fig. S2a and b). For murine mNLRP1b inflammasome, mNLRP1b can directly activate downstream signal pathways with or without ASC bridging, but the participation of ASC strongly promotes the activation level of mNLRP1b inflammasome (Fig. S2a and b). Moreover, the ASC-containing inflammasome assembly can be detected in 293T-ASC-GFP cells with mNLRP1b and NLRP1 expression, but not in CARD8-T48 and CARD8-T60 expressing cells (Fig. S2c and d). Our results are consistent with previous reports^20–23^ that ASC is an indispensable, dispensable and independent adaptor in NLRP1, mNLRP1b and CARD8 inflammasome assembly, respectively.

Because the C-terminal structural segment containing a UPA subdomain and an entire CARD domain is responsible to recruit downstream signal molecules to complete the inflammasome assembly when NLRP1, mNLRP1b and CARD8 are activated by danger signals^24^. We next examined whether the distinct ASC preference for NLRP1, mNLRP1b and CARD8 inflammasome depends on UPA subdomain or CARD domain. We first employed the mammalian two-hybrid (M2H) system to assess the interaction of NLRP1, mNLRP1b and CARD8 with ASC. The UPA-CARD fragment of NLRP1 and mNLRP1b, but not CARD8, showed a strong binding with ASC in agreement with the functional study results (Fig. 4a). In addition, the M2H results also showed that the homotypic CARD-CARD interaction plays an important role in mediating the interaction of NLRP1 and mNLRP1b with ASC (Fig. 4a). Then, we generated a series of chimeric proteins for the UPA-CARD domain of NLRP1, mNLRP1b and CARD8 (Fig. 4b). Chimera-1 and chimera-2, in which the NLRP1^CARD^ domain was substituted with the CARD domain of mNLRP1b or CARD8, could activate the inflammasome-mediated signals independent of ASC (Fig. 4c and 4d). Meanwhile, replacing the mNLRP1b^CARD^ or CARD8^CARD^ domain with CARD domain of NLRP1 (chimera-3 and chimera-4, respectively) blocked the pro-IL-1β maturation and secretion in the absence of ASC (Fig. 4c and 4d). By the analysis of the ASC-dependent sensor proteins, all NLRP1^UPA-CARD^, chimera-3 and chimera-4 share the common structural feature including the same CARD domain of human NLRP1. We conclude that the distinct ASC preference depends on the CARD domain in NLRP1, mNLRP1b and CARD8 inflammasome, while the UPA subdomain does not determine the specificity to assemble distinct inflammasome complexes.

**Fig. 4.**
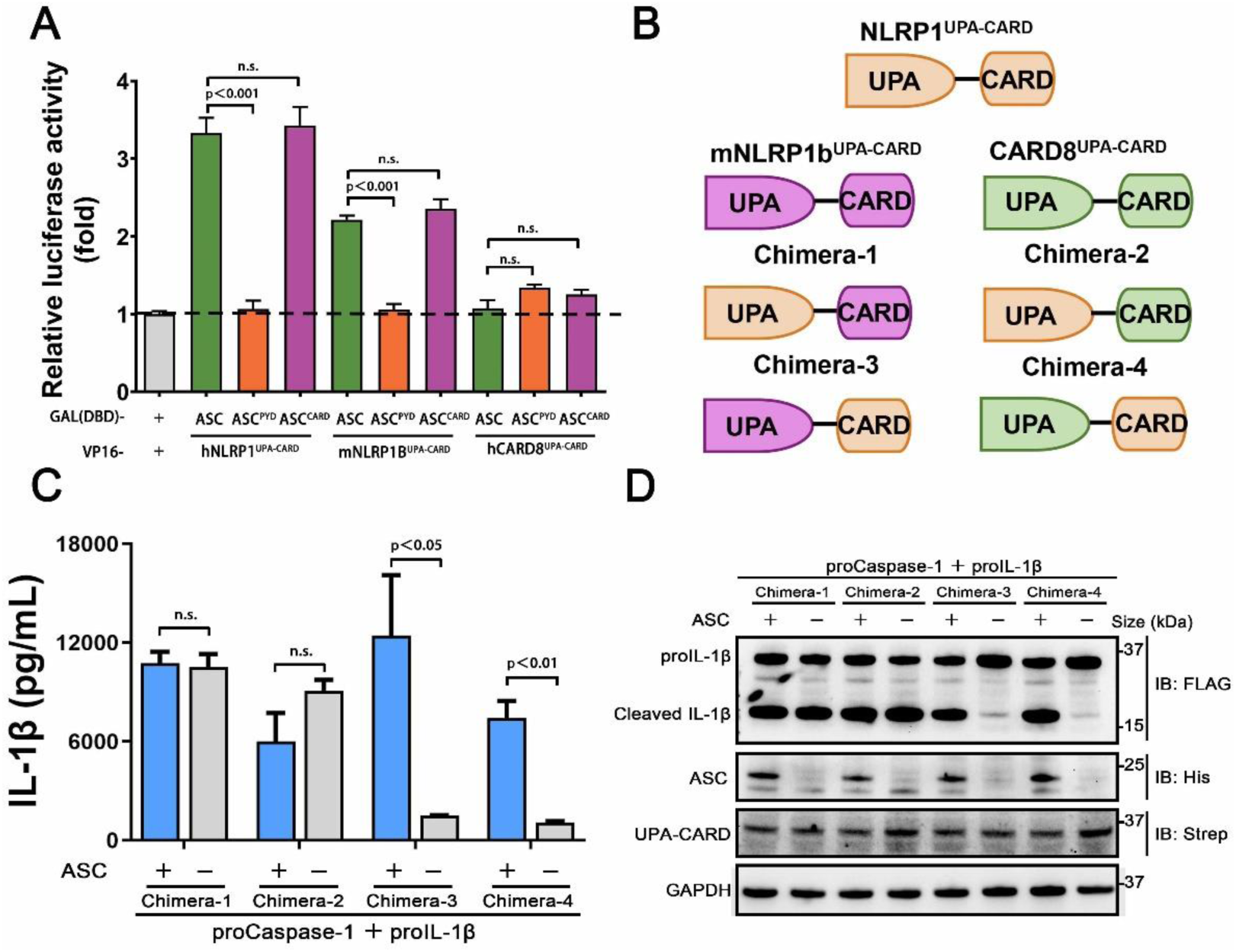
CARDs play the key roles for ASC dependence in NLRP1 and CARD8 inflammasome. **a** Mammalian two-hybrid characterization for the interaction of ASC with NLRP1^UPA-CARD^, mNLRP1b^UPA-CARD^ and CARD8^UPA-CARD^. Luciferase activity was normalized to *Renilla* and data were shown as the fold of empty vector control. Mean values ± SEM are representative of three biological replicates. **b** Diagram of the chimera proteins of NLRP1^UPA-CARD^, mNLRP1b^UPA-CARD^ and CARD8^UPA-CARD^. **c,d** ELISA and immunoblotting analysis were conducted from ASC-deficient chimera inflammasome. Data are shown as mean ± SEM and representative of three biological replicates.

### Structure basis for CARD domain-mediated ASC preference

According to the above results, CARD domain acts as a key role in the ASC dependence for the assembly of murine mNLRP1b, human NLRP1 and CARD8 inflammasome. Due to the close correlation between structure of proteins and molecular function, we next attempted to explain the inflammasomes assembly mechanism by three-dimensional structure analysis. The crystallographic structures of C-terminal CARD domain of NLRP1 and CARD8 were reported by prior studies^25, 26^, but the three-dimensional structure of mNLRP1b^CARD^ remains unknown. The predicted structure of mNLRP1b^CARD^ by AlphaFold has a high confidence because of conserved three-dimensional structure of Death Domain superfamily members^27, 28^ (Fig. 5a). The C-terminal CARD domain of the predicted mNLRP1b structure is composed of six anti-parallel α-helices folded in a Greek key arrangement (Fig. 5b). The five DD superfamily members, including mNLRP1b^CARD^, NLRP1^CARD^ (4IFP), CARD8^CARD^ (4IKM), CASP1^CARD^ (5FNA), ASC^CARD^ (6KI0), can be aligned with rmsds between 0.6 Å and 2.2 Å to confirm the conserved structural characteristics of CARD domains (Fig. 5c).

**Fig. 5.**
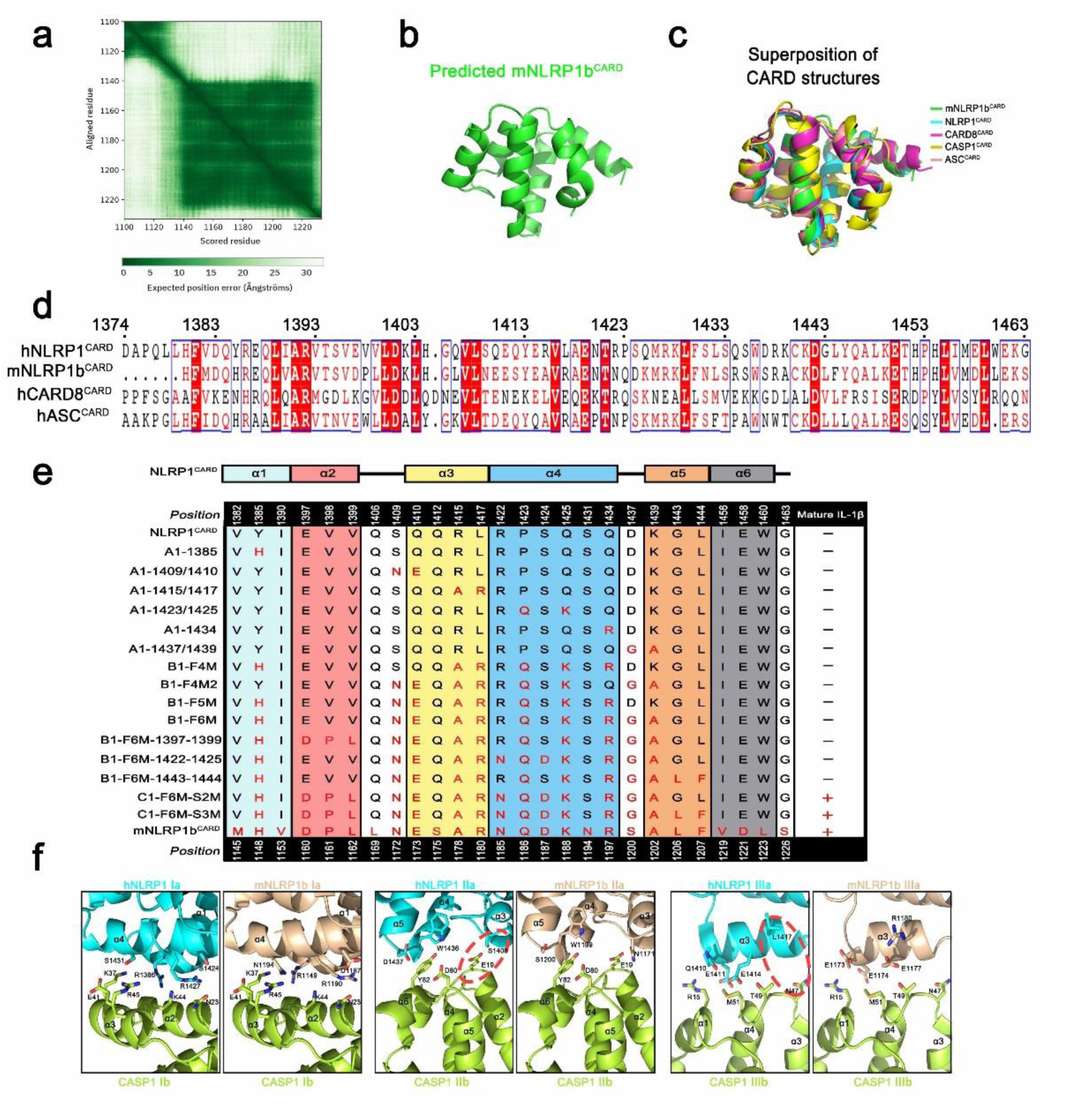
Various CARD interfaces were critical for ASC dependence in NLRP1 and CARD8 inflammasome. **a** The AlphaFold’s predicted aligned error of mNLRP1b encoding residues 1100-1233 structure. **b** Predicted structure of the CARD domain of mNLRP1b. **c** Superposition of all five known CARDs from mNLRP1b (green), NLRP1 (cyan), CARD8 (magenta), CASP1 (yellow), and ASC (lightpink). **d** Sequence alignment analysis of hNLRP1^CARD^, mNLRP1b^CARD^, CARD8^CARD^ and hASC^CARD^. Amino acid position is shown on the top of the sequences. **e** Mapping critical hNLRP1^CARD^ interaction sites for ASC-deficient NLRP1 inflammasome activation based on mNLRP1b^CARD^ sequence. The secondary structure of hNLRP1^CARD^ is indicated on the top. **f** Structural compatibility for Type Ⅰ, Ⅱ, and Ⅲ interaction residues of CASP1^CARD^ with human NLRP1^CARD^ or murine mNLRP1b^CARD^. Red line is used to label possible clashes in detailed CARD-CARD interaction model.

Since the structures of mNLRP1b^CARD^, NLRP1^CARD^ and CARD8^CARD^ showed a high similarity, we further determined the contribution of different amino acids in the CARD domain that may result in the distinct ASC preference. Sequence alignment of NLRP1^CARD^, mNLRP1b^CARD^, CARD8^CARD^ and ASC^CARD^ suggested a low sequence identity of CARD domains (Fig. 5d). Thus, we supposed that different amino acids in the CARD domain contribute to the ASC dependence for the inflammasome assembly. To prove this hypothesis, site-directed mutagenesis on NLRP1^UPA-CARD^ was conducted to map critical surface sites mediating the interaction between NLRP1^UPA-CARD^ and downstream signal molecule pro-Caspase-1 according to the primary sequence of mNLRP1b^UPA-CARD^ (Table. S1). The systematic site-directed mutagenesis analysis confirmed that the sixth α-helix of NLRP1^CARD^ does not affect NLRP1 inflammasome assembly to carry out signal amplification. More importantly, we also found that multiple molecular surfaces were involved in the interaction of NLRP1^UPA-CARD^ and Caspase-1^CARD^ to transduce downstream signals, which is consistent with the Mosaic model that DD superfamily members co-assemble into archetypical supramolecular platform by three conserved interaction types^21^ (Fig. 5e and S3a-d).

Type Ⅰ, Type Ⅱ, and Type Ⅲ interaction types of DD superfamily members, first defined in MyDDosome complex assembly, were widely adopted in describing supramolecular filament formations of DD superfamily^29^. On the other hand, the structural conformations of NLRP1^CARD^ and mNLRP1b^CARD^ are similar to the structure of ASC^CARD^ with rmsds 0.814 Å and 0.718 Å, respectively. To elucidate the heterotypic interactions between CARDs of sensors and downstream effector Caspase-1^CARD^, we modeled three asymmetric interfaces of DD superfamily between NLRP1^CARD^ or mNLRP1b^CARD^ and Caspase-1^CARD^ based on the cryo-EM structure of octamer with 4 ASC^CARD^ molecules and 4 Caspase-1^CARD^ molecules (PDB: 7KEU)^30^. As shown in Figure 5f, the interaction of mNLRP1b^CARD^ and Caspase-1^CARD^ is favored, whereas NLRP1^CARD^ clashes with Caspase-1^CARD^. For example, the E19 and N47 on the Caspase-1^CARD^ Type Ⅱb and Ⅲb surfaces show strong compatibility with corresponding positions N1171 and R1180 in mNLRP1b^CARD^, but not the S1409 and L1417 on the NLRP1^CARD^ Type Ⅱa and Ⅲa surfaces. Taken together, our results provide a detailed structural foundation for ASC preference during most CARD-containing inflammasome assembly.

### The functional implications of patient-derived NLRP1 mutations

With the development of whole-exome sequencing technology and further studies of NLRP1 related diseases, an increasing number of mutations of NLRP1 was reported in patients experienced different auto-inflammatory diseases^24, 31^. By summarizing all disease-causing mutations in NLRP1 and mapping NLRP1 mutations onto the predicted full-length structure of NLRP1, we found that these mutations are mainly located in three distinct molecular regions (Fig. 6a). The auto-inhibitory PYD domain of NLRP1 is the first region that includes A54T, A66V and M77T mutations resulting in rare monogenic inflammatory skin diseases (Fig. S4a). The second one is localized in the linker region between NACHT domain and LRR domain, including R726W, T755N and F787_R843del mutations (Fig. S4b). However, the molecular mechanism of how these mutations cause increased NLRP1 activation and autoimmune-inflammatory disease are unknown. M1184V and R1214R mutations located in the third region may affect the auto-processing of FIIND domain to release the C-terminal autoproteolytic fragment of NLRP1 (Fig. S4c).

**Fig. 6.**
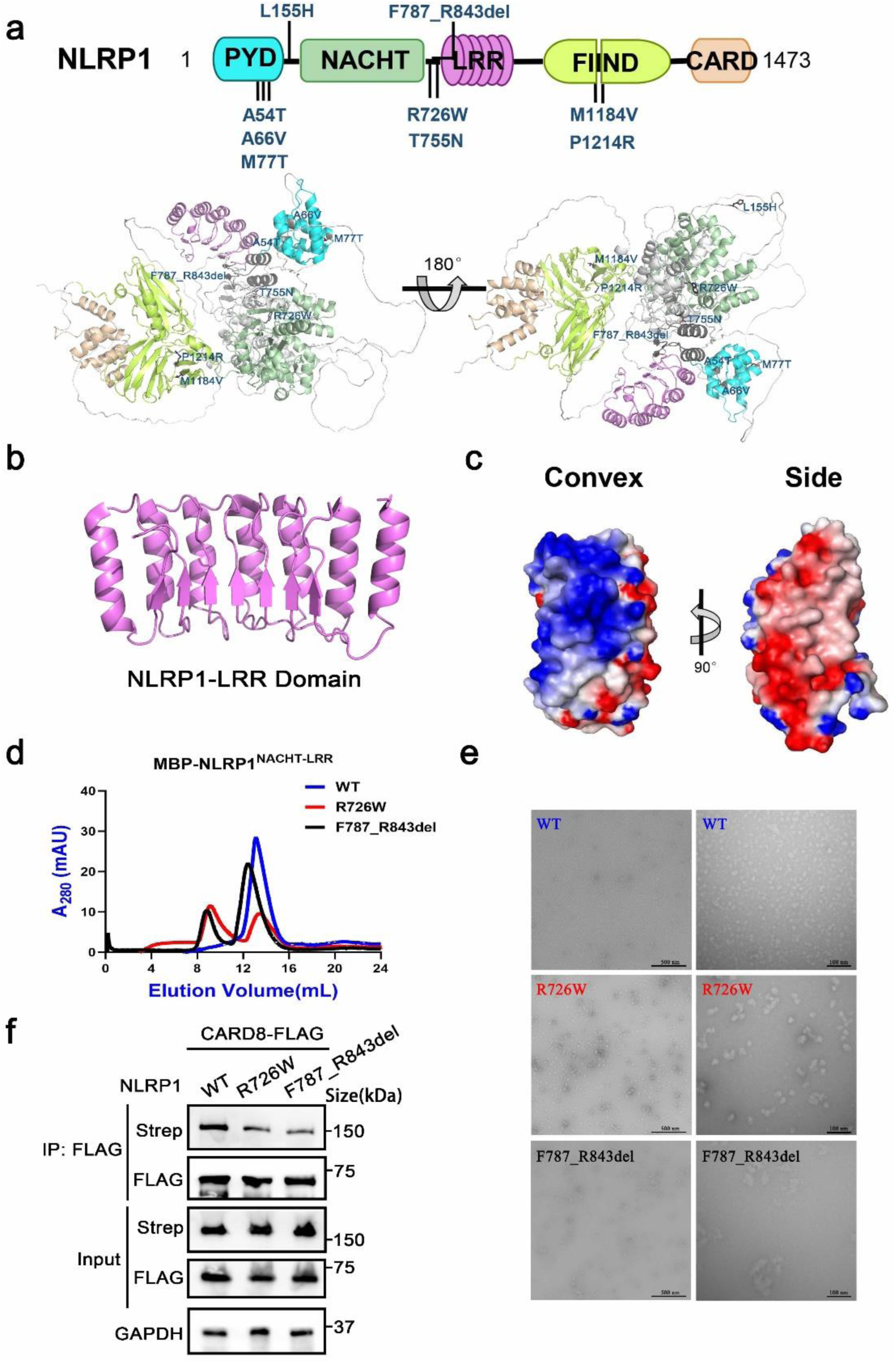
The disease-causing mutations cause decreased interaction between NLRP1 and CARD8. **a** The disease-causing mutations in autoimmune-inflammatory diseases are indicated on the predicted structure of full-length NLRP1. **b** Monomer crystal structure of LRR domain of NLRP1. **c** The electrostatic charge surfaces analysis of NLRP1^LRR^. The electrostatic potential of the convex and side of NLRP1^LRR^ was calculated by APBS. **d** Gel filtration characterization of recombinant MBP-NLRP1^NACHT-LRR^, R726W mutant and F787_R843del mutant proteins. **e** Representative negative-stain electron microscopy snapshots of recombinant MBP-NLRP1^NACHT-LRR^, R726W mutant and F787_R843del mutant proteins. **f** HEK 293T cells were transiently transfected with constructs for NLRP1, R726W mutant, F787_R843del mutant and CARD8. After 28 hours, the protein expression was detected by immunoblotting, and co-immunoprecipitation were conducted with anti-FLAG antibody to analyze the interaction.

Because the NACHT and LRR domains of NLRP1 are critical for the interaction with CARD8 and associated with several auto-inflammatory diseases. We next sought to study the structure of the NACHT and LRR domains of NLRP1 to elucidate the self-inhibition mechanism of NLRP1. The crystallographic structure of NLRP1^LRR^ revealed a compact structure of crescent shape in 2.45 Å resolution, which is composed of 7 parallel α helices and 7 parallel β strands (Fig. 6b and Table. S2). The NLRP1^LRR^ structure is nearly identical to the reported structure^32^. Electrostatic surface charge analysis of the NLRP1^LRR^ showed a polar surface pattern, in which the majority of the convex surfaces formed by the α-helices are heavily positively charged, while the surfaces formed by part of the convex and one side are heavily negatively charged (Fig. 6c). This result is consistent that NLRP1 could directly bind dsRNA via its LRR domain to trigger NLRP1 inflammasome activation^4^. On the other hand, the biochemical properties of R726W and F787_R843del mutations of NLRP1, located in the linker region between NACHT domain and LRR domain, were characterized by size-exclusion chromatography and the negative-stain EM analysis. As expected, the two disease-causing mutations of NLRP1 (R726W and F787_R843del) drastically increase oligomer formation as compared to wild-type NLRP1 NACHT-LRR domain, indicating that these two amino acid positions play significant roles in the auto-inhibition of human NLRP1 (Fig. 6d and 6e). More importantly, we note that R726W and F787_R843del mutations, which cause autoimmune and auto-inflammatory diseases, showed a considerably decreased binding with CARD8 (Fig. 6f). These findings suggest that the auto-inhibitory domains of NLRP1 are likely to maintain the native inactive monomer state of NLRP1 by interacting with CARD8.

## Discussion

The NLRP1 inflammasome, a macromolecular complex platform centered with NLRP1 as a sensor protein, has mainly made two remarkable progresses in recent years. First, it was reported that NLRP1 is activated by two physiological activators, including viral 3C proteases^3^ and double-stranded RNA^4^. Second, several recent studies have implied a role for NLRP1 mutations in multiple autoimmune diseases^31, 33^ and the pathogenic mechanistic basis of many disease-causing mutations was partially explained. Although human NLRP1, unlike other inflammasome sensors, has a wide tissue expression profile which suggests that it may play essential roles in both host defense and human diseases, the overall role of NLRP1 inflammasome in human remains elusive resulting from the lack of knowledge of its regulatory mechanisms.

The CARD8 gene is capable to generate different isoforms that bear distinct regulation functions in inflammatory signal transduction mechanism. The T60 isoform of CARD8, but not the T48 isoform, inhibits NLRP3 inflammasome activity by binding with NLRP3 and mediates the effect of the V44I mutation on NLRP3 functional activation^15^. Consistently, CARD8 was shown to suppress activation of NLRP3 through the CARD8-NLRP3 interaction and lost the inhibition ability because of NLRP3 with mutations associated with cryopyrin-associated periodic syndromes (CAPS)^16^. Moreover, a recent study proposed that CARD8 disturbs the formation of the NODosome complex via an attenuation of the NOD2-mediated oligomerization^14^. On the other hand, it is previously reported that DPP8/9 is an interacting partner of NLRP1 and maintains NLRP1 in its inactive state to avoid NLRP1 inflammasome downstream signal transduction^8^. Our results identified CARD8 as a novel inflammasome regulator for human NLRP1 and provided a structural basis for the specificity of signaling pathways in ASC-indispensable, ASC-dispensable and ASC-independent inflammasomes.

In this study, we identified that the “canonical” T48 isoform of CARD8 interacts with human inflammasome sensor NLRP1, but fails to suppress its activation. Surprisingly, we found the somewhat larger T60 isoform of CARD8 could act as a negative regulator to control the NLRP1 inflammasome activation. Mechanistically, *in vitro* binding assay revealed CARD8-T60 isoform and NLRP1 directly bind to each other. Using a series of truncation mutants of CARD8-T60 isoform and NLRP1, we found that CARD8-T60 variant interacts with NLRP1 through ZU5 subdomain, while NLRP1 interacts with CARD8 through the NACHT, LRR and FIIND domains. In addition, we demonstrated that ASC preference of CARD-containing inflammasomes results from the CARD domain of these receptors, but not the UPA subdomain. Different amino acids in the CARD domain are major contributors of ASC preference and the sixth α-helix of CARD domain does not mediate the hetero-oligomerization in signaling transduction event. Finally, our findings also indicated the patient-derived mutations lead to the NLRP1 oligomerization and disrupt the NLRP1-CARD8 interaction, thereby providing a mechanistic explanation for auto-inflammatory diseases. In summary, our cellular, biochemical and structural evidence lead us to propose the NLRP1 and CARD8 functional model (Fig. 7). In resting state, the auto-inhibitory function of wild-type full-length NLRP1 is maintained by NLRP1 and CARD8 interaction. While these patient-derived NLRP1 mutations could disrupt the self-inhibitory mechanism of NLRP1 by weakening the binding of NLRP1 and CARD8. This triggers constitutive self-oligomerization of NLRP1 and CARD8 and recruits ASC or CASP1 to assemble the distinct inflammasomes which is participated by multiple molecular surfaces of CARDs.

**Fig. 7.**
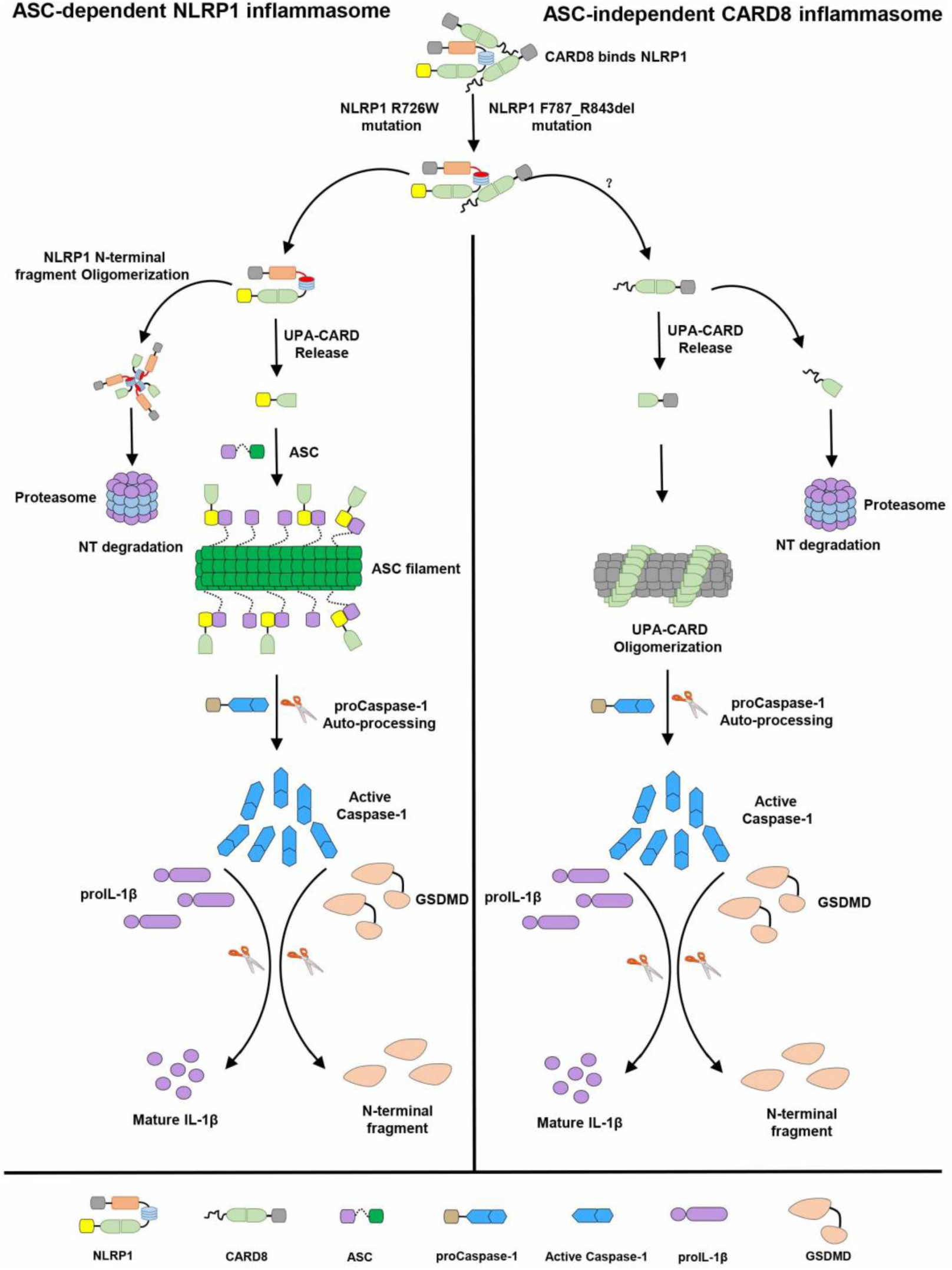
The sequential regulation and activation model of NLRP1 and CARD8 inflammasome. Full-length NLRP1 interacts with CARD8 to maintain the inactive state of NLRP1 and CARD8 in resting cells. The disease-causing mutations would lead to decrease the interaction between NLRP1 and CARD8 and lost the auto-inhibitory mechanism of NLRP1. Then NLRP1 and CARD8 N-terminal degradation by proteasome results in the release of the active C-terminal segment (UPA-CARD domain) from NLRP1 and CARD8. Oligomerized UPA-CARD fragments differentially recruit ASC or CASP1 to assemble the distinct inflammasomes by multiple molecular surfaces of CARDs.

The aberrant activation of NLRP1 inflammasome has been implicated in many auto-inflammatory diseases and the common treatment strategy of these diseases is to block the IL-1β maturation and secretion^34, 35^. But NLRP1 inflammasome also results in cleavage of pro-IL-18 and pyroptosis, which are most pathogenic and pro-inflammatory. A useful therapy for treating these diseases is to target the upstream signaling molecules to disturb NLRP1 inflammasome activation. Thus, CARD8 and NLRP1 interaction identified by our results might provide a therapeutic target for the treatment of NLRP1-mediated diseases.

## Materials and Methods

### Cloning and Reagents

Constructs for entire coding sequence of ASC, proCaspase-1 and IL-1β were prepared as previously described^21^. The cDNA sequences of mNLRP1b (UniProt ID: Q2LKW6-1), NLRP1 (UniProt ID: Q9C000-1), CARD8-T48 (UniProt ID: Q9Y2G2-1) and CARD8-T60 (UniProt ID: Q9Y2G2-5) were amplified from mouse or human spleen cDNA libraries and their truncations were constructed by PCR based on the full-length genes. The QuikChange technology was used to generate point mutations of NLRP1 and CARD8. The constructs of UPA-CARD fragment of NLRP1, mNLRP1b and CARD8 were cloned into pACT vector with VP16 activation domain, whereas DNA encoding full-length ASC and truncated ASC were shuttled into pBIND vector with DNA-binding domain and *Renilla reniformis* luciferase sequence. The constructs containing RFP or GFP were used for fluorescence co-localization assay. Other plasmids for expression in HEK 293T cells were inserted into modified pcDNA3.1 vectors using standard cloning techniques. The NLRP1^LRR^ (V790-S990) and NLRP1^NACHT-LRR^ (G230-P994) were engineered into a pET30a-derived vector with a Tobacco Etch Virus (TEV) protease linker between N-terminal fusion tag and target protein via homologous recombination. All plasmids were confirmed by Sanger sequencing. The reagents used include Human IL-1β ELISA kit (BD Biosciences, 557953), anti-FLAG antibody (Sigma, F1804), anti-Strep (Biolegend, 688202), anti-GAPDH (ABclonal, AC002), anti-His (Sangon Biotech, D191001), anti-GFP (Abmart, M20004M), FLAG Beads (Sigma, A2220), Strep Beads (IBA, 2-1208-025), Z-VAD-FMK (APExBio, A1902), Val-boroPro (TargetMol, T4042), Disuccinimidyl suberate (DSS) (PIERCE, 21555).

### Cell culture and stable cell lines generation

HEK 293T cells were purchased from Cell Bank of Chinese Academy of Sciences and grown in Dulbecco’s modified Eagle’s medium (DMEM) supplemented with 10% FBS, 100 U/ml penicillin and 100 μg/ml streptomycin. Transfection of HEK 293T with 2 μg of ASC-GFP-pCDH, 2 μg psPAX2 and 2 μg pMD2.G was performed to package the lentivirus. The virus supernatants with 10 μg/ml polybrene were used to infect HEK 293T cells. 36 hours later, the cells were selected for ectopically expressing ASC-GFP HEK 293T cell line using 5 μg/ml puromycin. All cells were cultured at 37 ℃ in an incubator containing 5% CO_2_.

### Determination of cleaved IL-1β of NLRP1 inflammasome activation assay

HEK 293T cells were plated at 1×10^5^ per well in twenty-four wells plates overnight before transfection. Then each well of cells was co-transfected with NLRP1, ASC, Caspase-1 and IL-1β plasmids to constitute NLRP1 inflammasome using lipo6000^TM^ at a “20:10:2:35” ratio according to the instructions of manufacturer. 24 hours after transfection, the media was changed with fresh medium for another 24 hours. The concentrations of IL-1β in cell lysates were measured by immunoblotting. Determination of secreted IL-1β in cell culture supernatants was performed with ELISA kits (BD Biosciences) according to standard manufacturer’s instructions without modification.

### SDS-PAGE, Immunoblotting and Immunoprecipitation

Cells were treated with ice-cold lysis buffer (20 mM Tris-HCl pH 7.5, 100 mM KCl, 5 mM MgCl_2_, 0.3% NP-40 and protease inhibitor cocktail) for 30 mins and centrifuged at 12000 rpm for 15 mins. The soluble fraction was incubated with Anti-FLAG M2 agarose beads or Strep-Tactin superflow affinity gel overnight at 4 ℃. After the gels were washed three times with lysis buffer, the bound proteins were eluted by boiling, separated by 15% SDS-PAGE, and visualized by Coomassie blue staining or immunoblotting. For immunoblotting, the PVDF membranes were used to transfer proteins, blocked by 5% non-fat milk, and incubated with respective antibodies at 4 ℃. The protein bands were developed with ECL reagents and visualized by Bio-Rad Gel Doc System.

### ASC crosslinking

In ASC crosslinking assay, cells were lysed with cell lysis buffer, and cell lysates were centrifuged at 2500g at 4 ℃ for 10 min. Cell pellet fractions were resuspended and washed for three times with PBS. The washed pellets were cross-linked for 30 min at room temperature with 2 mM DSS. After centrifugation at 15000 rpm for 10 min, the samples were then combined with SDS loading buffer, heated to 100 ℃ for 10 min and carried out for immunoblot analysis.

### Fluorescence co-localization and ASC-GFP speck quantification assay

A density of 5×10^5^ cells was seeded on glass-bottom dishes, followed by fixedness with 4% paraformaldehyde 26 hours post transfection as described. Live-cells were incubated with 1 μg/ml DAPI for the sake of nucleus staining and visualized using a fluorescence microscope (OLYMPUS, TH4-200). For ASC-GFP speck quantification assay, total cell area was counted using blue fluorescence channel images. Quantification of ASC Speck was performed by the “Analyze particles” function following threshold adjustment based on GFP-positive signal in ImageJ/FlJl software.

### Mammalian two-hybrid assay

Based on Checkmate mammalian two-hybrid system (Promega, Madison, WI, USA), the indicated genes were engineered into pACT and pBIND vectors, respectively. These pACT, pBIND and pG5*luc* plasmids at 1:1:1 molar ration were co-transfected into HEK 293T cells following the manufacturer’s manual. The relative luciferase activity was determined by Dual Luciferase Reporter Gene Assay Kit (Beyotime) after 24 hours post transfection. The empty plasmids were also transfected into cells as a negative control and three independent experiments were performed for data acquisition.

### Recombinant protein preparation

Recombinant NLRP1^LRR^ tagged with a removable GB-1 was expressed following Studier and colleagues’ protocol^36^. The BL-21-CodonPlus RIPL strain (Strategen) was transformed with expression plasmids encoding NLRP1^LRR^ protein. Bacteria were grown in LB medium and induced with 0.3 mM isopropyl 1-thio-β-D-galactopyranoside (IPTG) when OD600 reached 0.6-0.8 at 16 ℃. The bacterial samples were harvested by centrifugation at 15000 rpm for 25 min and lysed by sonication, followed by Ni-NTA chromatography. The recombinant GB-1-NLRP1^LRR^ protein was added with TEV protease to remove the GB-1 tag and separated by the second step of Ni-NTA purification. The non-tagged NLRP1^LRR^ protein was further purified from eluant to homogeneity by size exclusion chromatography with a soluble buffer containing 100 mM NaCl, 5% glycerol, and 20 mM HEPES pH 7.4.

Expression and purification of NLRP1^NACHT-LRR^ and its mutated proteins were similar to the above. Following sonication and centrifugation in lysis buffer (50 mM Tris-HCl at pH 7.5, 10% glycerol, 500 mM NaCl, 20 mM imidazole and 0.01% NP40), the clarified supernatant containing the NLRP1^NACHT-LRR^ protein with maltose-binding protein (MBP) tag was purified on Hisprep IMAC column (GE Healthcare) and eluted with elution buffer contained 50 mM Tris-HCl at pH 7.5, 10% glycerol, 500 mM NaCl, 300 mM imidazole and 0.01% NP40. To further purify the final MBP-NLRP1^NACHT-LRR^ protein, size exclusion chromatography was performed with an XK26/60 Superdex 200 size exclusion column in 50 mM Tris-HCl at pH 7.5, 10% glycerol, 300 mM NaCl and 0.01% NP40 buffer.

### Negative-stain EM

In brief, the carbon coated 400-mesh Cu EM specimen grids (Solarus, Gatan, Model 950) were glow discharged before 50-100 μg/ml of protein samples were left on the grids for 90 s. After the samples were blotted, stain containing 2% Uranyl Acetate was employed to incubate with the grids for 60s. Following air dry of the grids, negative stained images were taken by 120 keV electron microscopy.

### Size-exclusion chromatography assay

To assess the oligomeric state of NLRP1^NACHT-LRR^ and its mutated proteins, the protein samples were subjected to analytical gel filtration using an XK26/60 Superdex 200 size exclusion column equilibrated with a buffer containing 50 mM Tris-HCl at pH 7.5, 10% glycerol, 300 mM NaCl and 0.01% NP40.

### Crystallography

The ∼50 mg/ml NLRP1^LRR^ protein was mixed at a 1:1 ratio with reservoir solution for crystallization conditions screening. Diffraction quality crystals were obtained by hanging-drop vapor-diffusion method with reservoir condition of 1.6 M Sodium Potassium Phosphate, 100 mM HEPES, pH 6.4. Typical diamond shaped crystals of 0.3-0.5 µm in each dimension grow in 2 weeks at room temperature. A final concentration of 20% glycerol was added into the crystallization condition as a cryoprotectant solution before flash-cooled the crystals in liquid nitrogen for data collection. X-ray diffraction data were collected at beamline X12C, X29A and X25 at the National Light Source, the Brookhaven National Laboratory and beamline SER-CAT 22ID and GM-CA 23ID at the Advanced Photon Source (APS), the Argonne National Laboratory. Data were processed with *HKL2000*^37^ and XDS^38^. In order to solve the phase problem, a homology structure model was built with the Swiss-Model server^39^ using the crystal structure of ribonuclease inhibitor (PDB: 1DFJ)^40^ as template. Molecular replacement was calculated in *Phaser*^41^ in the CCP4i GUI^42^ by using 4 central repeats of the LRR homology model as an initial search model. The structure model was completed by manual building in *Coot*^43^. Structure refinement was performed by REFMAC5^44^, CNS^45^ and PHENIX^46^. Structural figures were prepared by *Pymol*.

### Statistical analysis

GraphPad Prism 8.0 software was used for statistical analysis and error bars were plotted as mean±SEM. Statistically significant of three independent biological experiments was determined by student’s unpaired *t*-test, and *P*-value less than 0.05 was denoted significant.

### Accession number

The accession number for coordinates and structural factors of NLRP1^LRR^ in this study is 5Y3S. The data of this manuscript can be obtained from the corresponding author upon request.

## Supporting information

5y3s structure validation report

## Acknowledgements

This study is supported by Strategic Priority Research Program of the Chinese Academy of Sciences (Grant No. XDB29030104), the National Natural Science Foundation of China (Grant No. 31870731, 81872110 and 82172773).

## Author contributions

ZX designed, carried out almost all experiments and wrote the first draft of the paper. SD, YH, YY, DZ, and PZ collected, analyzed and interpreted the data. LS, WZ, XY, ML, HL, and WZ contributed to the helpful discussion and comment on experiment results. YZ and TJ provided study design, supervision and data analysis, and approved the final version of the manuscript.

## Compliance with ethical standards

### Conflict of interest

The authors declare no financial conflicts of interests exist.

## Notes

https://www.rcsb.org/structure/5Y3S

### Competing Interest Statement

The authors have declared no competing interest.

