## Supplemental Figures and Tables for "CARD8 negatively regulates NLRP1 inflammasome activation level by interaction with NLRP1"

**Supplementary Materials**

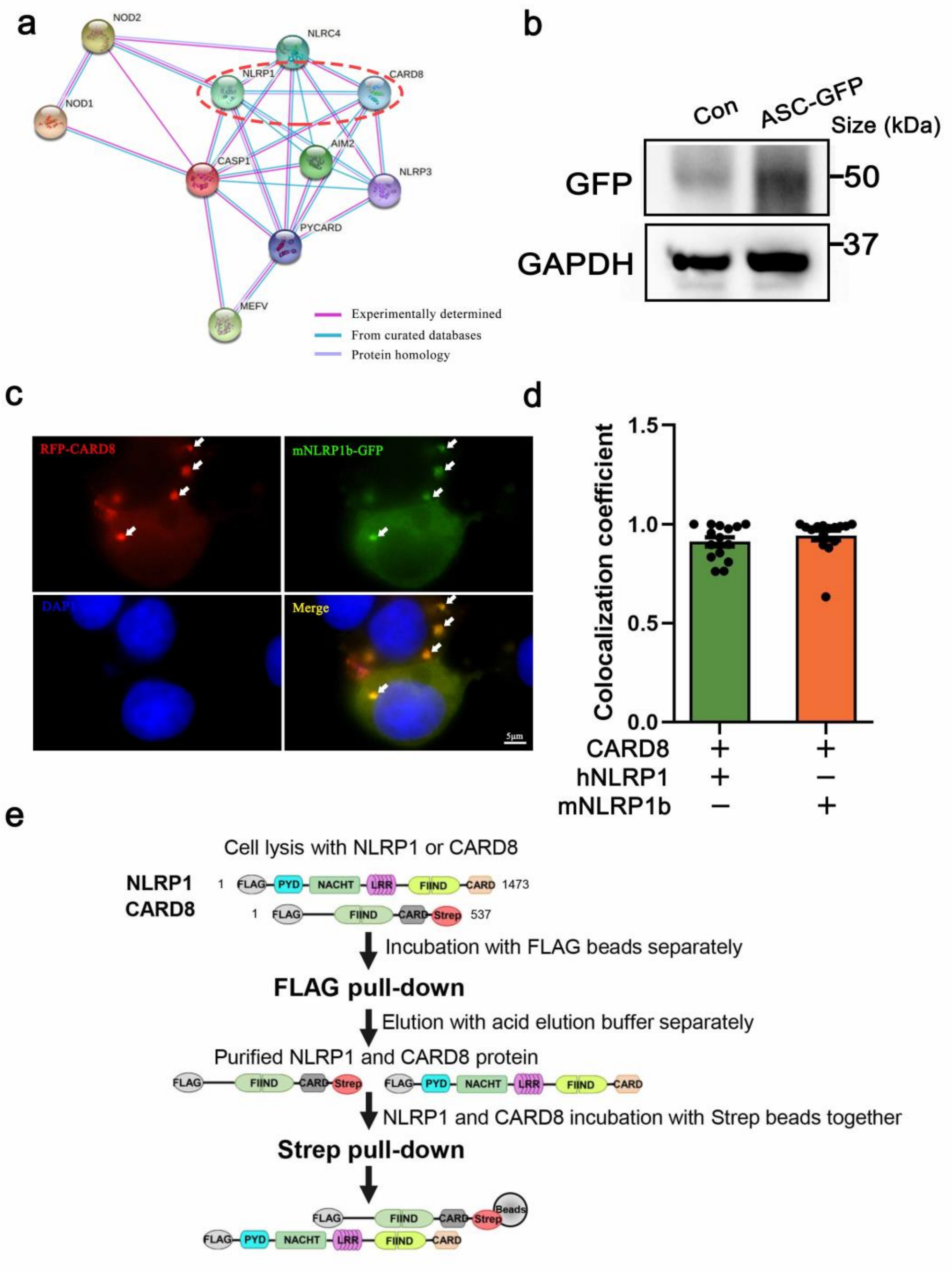


**Fig. S1** Physical interaction between NLRP1 and CARD8. **a** Protein-protein interaction network of the inflammasome scaffold molecules. Different line colors represent different types of protein associations. Experimental (red), Curated databases (light blue), Protein homology (purple). **b** Generation of the HEK 293T cell line stably expressing ASC-GFP. Efficiency of knock-in was evaluated using western blot. **c** Transient transfection RFP-CARD8 and mNLRP1b-GFP in HEK 293T cells. Green and red fluorescent were captured by fluorescence microscopy. Co-localization regions are labeled with white arrow. **d** Colocalization coefficient of CARD8 with NLRP1 or mNLRP1b in HEK 293T cells (n=15 cells from three biological replicates). **e** Schematic drawing of the pull-down strategy by anti-FLAG and anti-Strep antibody for detection of NLRP1 and CARD8 direct interaction.

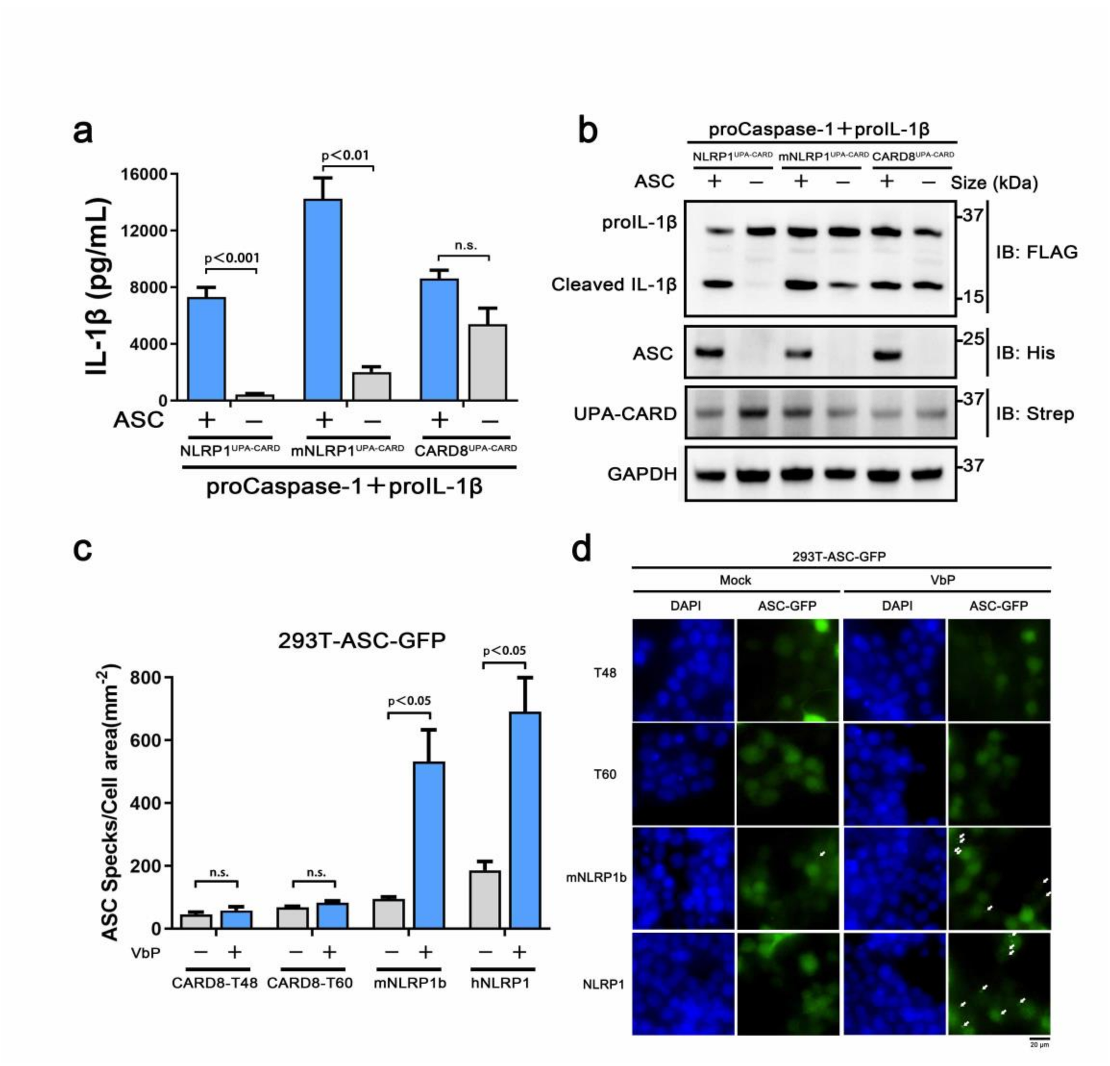


**Fig. S2** Distinct assembly mechanism for ASC dependence in NLRP1 and CARD8 inflammasome. **a,b** ELISA and immunoblotting analysis were conducted from ASC-deficient NLRP1, mNLRP1b and CARD8 inflammasome. Data are shown as mean ± SEM and representative of three biological replicates. **c,d** HEK 293T cells stably expressing ASC-GFP were transiently transfected with the indicated expression plasmids and treated with VbP (5 μM) for 6 hours. The cells were fixed with 4% formaldehyde and evaluated for ASC speck formation by fluorescence microscopy in three differential regions. Data of mean specks per cell area are shown as mean ± SEM in c and representative images are shown in d.

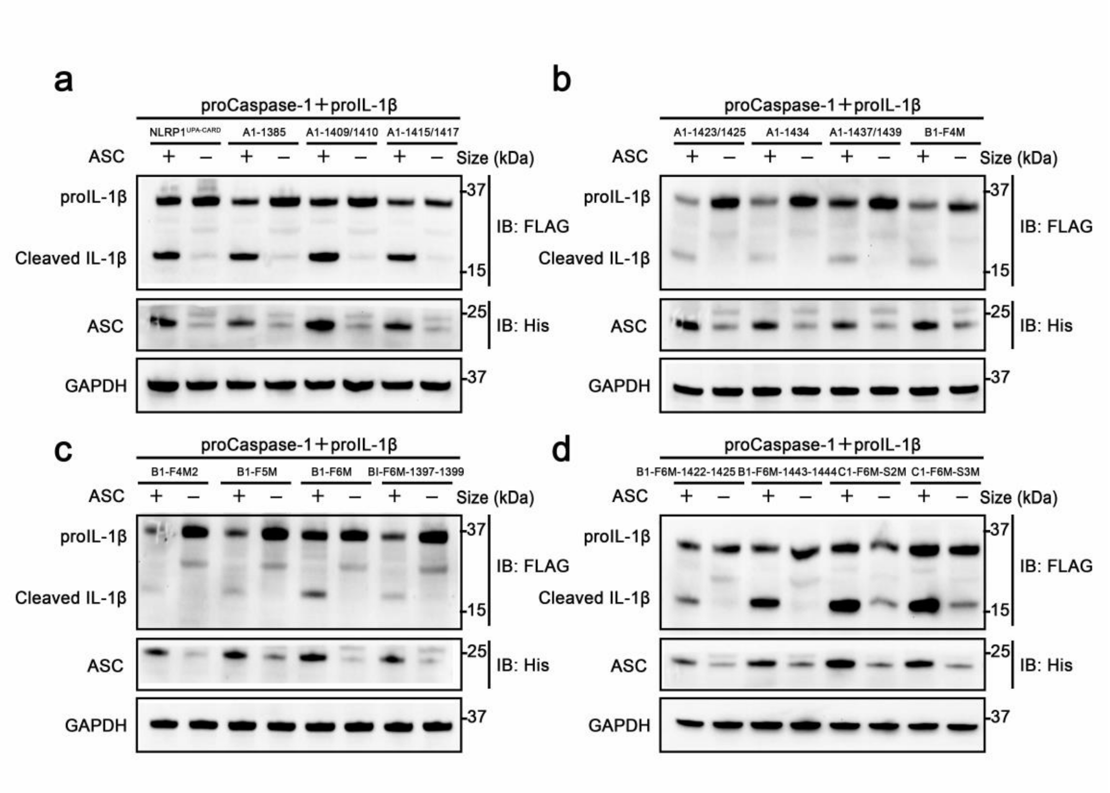


**Fig. S3** NLRP1 CARD domain mutations analysis using NLRP1 inflammasome reconstitution assay in HEK 293T cells. **a** Immunoblotting analysis were conducted from wild-type, A1-1385, A1-1409/1410 and A1-1415/1417 mutations of NLRP1$^{UPA\text{-}CARD}$ in reconstituted NLRP1 inflammasome system. **b** Immunoblotting analysis were conducted from A1-1423/1425, A1-1434, A1-1437/1439 and B1-F4M mutations of NLRP1$^{UPA\text{-}CARD}$ in reconstituted NLRP1 inflammasome system. **c** Immunoblotting analysis were conducted from B1-F4M2, B1-F5M, B1-F6M and B1-F6M-1397-1399 mutations of NLRP1$^{UPA\text{-}CARD}$ in reconstituted NLRP1 inflammasome system. **d** Immunoblotting analysis were conducted from B1-F6M-1422-1425, B1-F6M-1443-1444, C1-F6M-

S2M and C1-F6M-S3M mutations of NLRP1$^{UPA-CARD}$ in reconstituted NLRP1 inflammasome system.

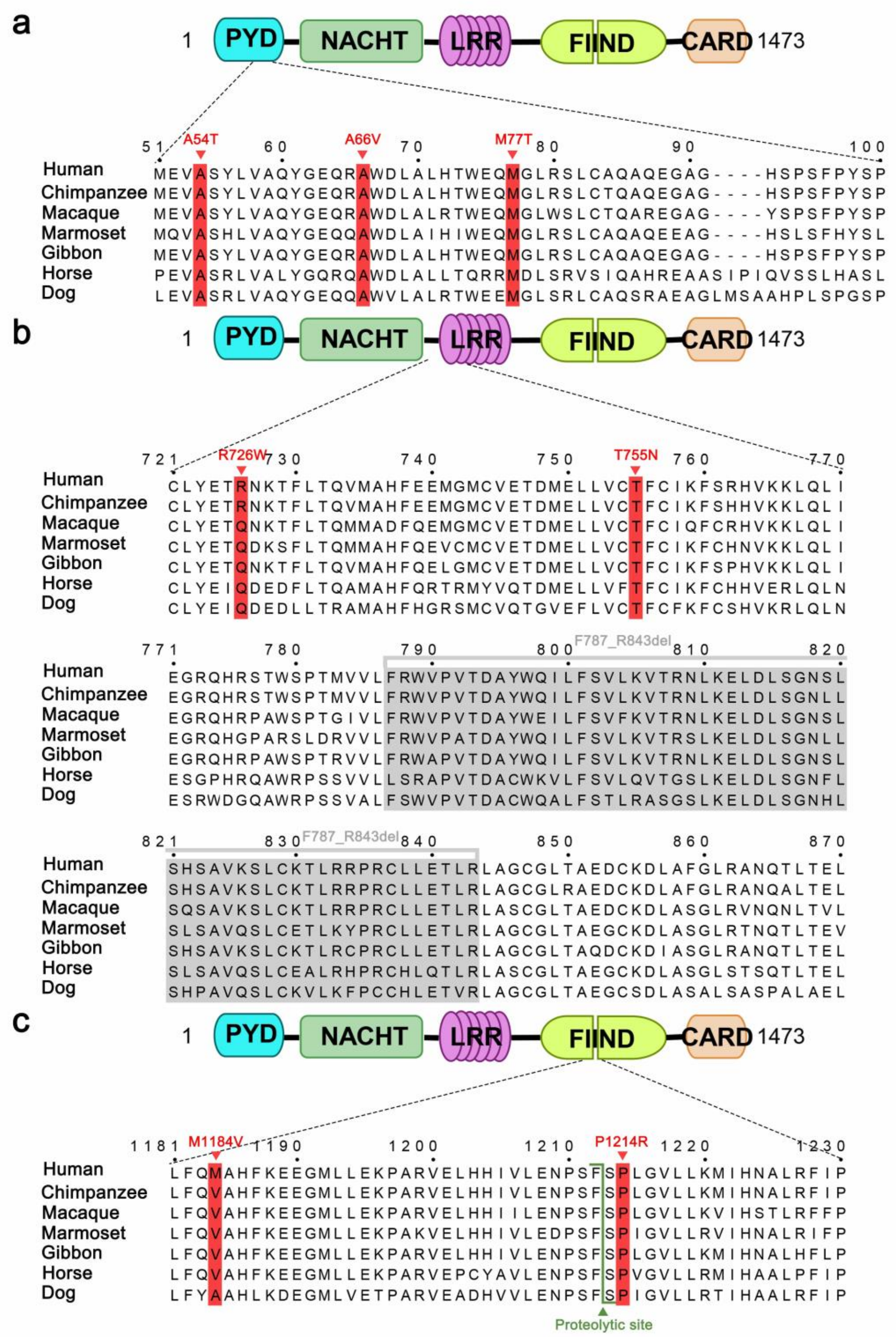

**Fig. S4** Sequence alignment analysis of NLRP1 from indicated species. **a** Sequence alignment analysis of PYD domain of NLRP1 from indicated species. Disease-causing mutations are highlighted in red. **b** Sequence alignment analysis of interdomain linker between NLRP1$^{NACHT}$ domain and NLRP1$^{LRR}$ domain from indicated species. Disease-causing mutations are highlighted in red or marked with gray shade. **c** Sequence alignment analysis of interdomain sequences between NLRP1$^{ZU5}$ subdomain and NLRP1$^{UPA}$ subdomain from indicated species. Disease-causing mutations are highlighted in red and proteolytic site is marked with green triangle arrow.

| Vectors and plasmids | Abbreviation | Mutated amino acid sites |
|---|---|---|
| pcDNA3.1 Strep-NLRP1 UPA-CARD | A1-1385 | Y1385H |
| pcDNA3.1 Strep-NLRP1 UPA-CARD | A1-1409/1410 | S1409N, Q1410E |
| pcDNA3.1 Strep-NLRP1 UPA-CARD | A1-1415/1417 | R1415A, L1417R |
| pcDNA3.1 Strep-NLRP1 UPA-CARD | A1-1423/1425 | P1423Q, Q1425K |
| pcDNA3.1 Strep-NLRP1 UPA-CARD | A1-1434 | Q1434R |
| pcDNA3.1 Strep-NLRP1 UPA-CARD | A1-1437/1439 | D1437G, K1439A |
| pcDNA3.1 Strep-NLRP1 UPA-CARD | B1-F4M | Y1385H,R1415A,L1417R,P1423Q,Q1425K,Q1434R |
| pcDNA3.1 Strep-NLRP1 UPA-CARD | B1-F4M2 | S1409N,Q1410E,R1415A,L1417R,P1423Q,Q1425K,D1437G,K1439A |
| pcDNA3.1 Strep-NLRP1 UPA-CARD | B1-F5M | Y1385H,S1409N,Q1410E,R1415A,L1417R,P1423Q,Q1425K,Q1434R |
| pcDNA3.1 Strep-NLRP1 UPA-CARD | B1-F6M | Y1385H,S1409N,Q1410E,R1415A,L1417R,P1423Q,Q1425K,Q1434R,D1437G, K1439A |
| pcDNA3.1 Strep-NLRP1 UPA-CARD | B1-F6M-1397-1399 | Y1385H,E1397D,V1398P,V1399L,S1409N, Q1410E, R1415A,L1417R,P1423Q,Q1425K,Q1434R,D1437G,K1439A |
| pcDNA3.1 Strep-NLRP1 UPA-CARD | B1-F6M-1422-1425 | Y1385H,S1409N,Q1410E,R1415A,L1417R,R1422N,P1423Q,S1424D,Q1425K, Q1434R,D1437G,K1439A |
| pcDNA3.1 Strep-NLRP1 UPA-CARD | B1-F6M-1443-1444 | Y1385H,S1409N,Q1410E,R1415A, L1417R,P1423Q,Q1425K,Q1434R,D1437G,K1439A,G1443L,L1444F |
| pcDNA3.1 Strep-NLRP1 UPA-CARD | C1-F6M-S2M | Y1385H, E1397D,V1398P, V1399L,S1409N,Q1410E,R1415A,L1417R,R1422N,P1423Q,S1424D,Q1425K,Q1434R,D1437G,K1439A |
| pcDNA3.1 Strep-NLRP1 UPA-CARD | C1-F6M-S3M | Y1385H,E1397D,V1398P, V1399L,S1409N, Q1410E,R1415A,L1417R,R1422N,P1423Q,S1424D,Q1425K,Q1434R,D1437G,K1439A,G1443L,L1444F |

**Table S2 X-ray Data Collection and Structure Refinement of NLRP1$^{LRR}$**

| | NLRP1$^{LRR}$ |
|---|---|
| **Data Collection** | |
| Space group | R32 |
| Unit cell (a, b, c) (Å) | 141.48, 141.48, 294.00 |
| (α, β, γ) (°) | 90, 90, 120 |
| Wavelength (Å) | 0.9000 |
| Wilson B-factor (Å$^2$) | 59.55 |
| Resolution (last shell) (Å) | 33.76 - 2.451 (2.539 - 2.451) |
| No of reflections (total/unique) | 223505 (22677) |
| Completeness (%) | 98.9 (99.1) |
| Average multiplicity | 5.4 (5.6) |
| Mean $I/\sigma(I)$ | 18.99 (1.90) |
| $R_{meas}$ (%) | 6.4 (93.9) |
| $R_{merge}$ (%) | 5.716 (95.33) |
| $R_{pim}$ (%) | 2.682 (43.63) |
| $CC_{1/2}$ | 0.999 (0.678) |
| **Refinement** | |
| Resolution (Å) | 50 - 2.45 |
| Reflections used in refinement | 41415 |
| No. of protein atoms /*B*-factor (Å$^2$) | 6210/70.45 |
| No. of heteroatoms/*B*-factor (Å$^2$) | 19/84.0 |
| RMSD bond lengths (Å) | 0.009 |
| RMSD bond angles (°) | 1.07 |
| $R_{work}$ (%)† | 21.25 |
| $R_{free}$ (%)‡ | 25.93 |

| | |
|---|---|
| Ramachandran plot favored/disallowed (%)** | 95/0 |
| PDB code | 5Y3S |

¶ $R_{merge} = \Sigma_h \Sigma_i |I_i(h) - \langle I(h)\rangle| / \Sigma_h \Sigma_i I_i(h)$, where $I_i(h)$ and $\langle I(h)\rangle$ are the ith and mean measurement of the intensity of reflection *h*.

¶ $R_{meas} = \Sigma_h(n/n-1)^{1/2}\Sigma_i |I_i(h) - \langle I(h)\rangle| / \Sigma_h \Sigma_i I_i(h)$, where $I_i(h)$ and $\langle I(h)\rangle$ are the *i*th and mean measurement of the intensity of reflection *h*.

† $R_{work} = \Sigma_h||F_{obs}(h)|-|F_{calc}(h)|| / \Sigma_h|F_{obs}(h)|$, where $F_{obs}(h)$ and $F_{calc}(h)$ are the observed and calculated structure factors, respectively. No $I/\sigma$ cutoff was applied.

‡ $R_{free}$ is the R value obtained for a test set of reflections consisting of a randomly selected 5% subset of the data set excluded from refinement.

**Values from Molprobity server (http://molprobity.biochem.duke.edu/).
